# Mosaic Turner syndrome shows reduced phenotypic penetrance in an adult population study compared to clinically ascertained cases

**DOI:** 10.1101/177659

**Authors:** Marcus A. Tuke, Katherine S. Ruth, Andrew R. Wood, Robin N. Beaumont, Jessica Tyrrell, Samuel E. Jones, Hanieh Yaghootkar, Claire L.S. Turner, Mollie E. Donohoe, Antonia M. Brooke, Morag N. Collinson, Rachel M. Freathy, Michael N. Weedon, Timothy M. Frayling, Anna Murray

## Abstract

Women with X chromosome aneuploidy such as 45,X (Turner syndrome) or 47,XXX (Triple X syndrome) present with characteristics including differences in stature, increased cardiovascular disease risk and primary ovarian insufficiency. Many women with X chromosome aneuploidy undergo lifetime clinical monitoring for possible complications. However, ascertainment of cases in the clinic may mean that the phenotypic penetrance is overestimated. Studies of prenatally ascertained X chromosome aneuploidy cases have limited follow-up data and so the long-term consequences into adulthood are often not reported. We aimed to characterise the prevalence and phenotypic consequences of X chromosome aneuploidy in a large population of women over 40 years of age. We detected 30 women with 45,X, 186 with mosaic 45,X/46,XX and 110 with 47,XXX among 244,848 UK Biobank women, using SNP array data. The prevalence of non-mosaic 45,X (1/8,162) and 47,XXX (1/2,226) was lower than expected, but was higher for mosaic 45,X/46,XX (1/1,316). The characteristics of women with 45,X were consistent with the characteristics of a clinically recognised Turner syndrome phenotype, including a 17.2cm shorter stature (SD = 5.72cm; *P* = 1.5 × 10^−53^) and 16/30 did not report an age at menarche. The phenotype of women with 47,XXX included taller stature (5.3cm; SD = 5.52cm; *P* = 5.8 × 10^−20^), earlier menopause age (5.12 years; SD = 5.1 years; *P* = 1.2 x 10^−14^) and a lower fluid intelligence score (24%; SD = 29.7%; *P* = 3.7 × 10^−8^). In contrast, the characteristics of women with mosaic 45,X/46,XX were much less pronounced than expected. Women with mosaic 45,X/46,XX were less short, had a normal reproductive lifespan and birth rate, and no reported cardiovascular complications. In conclusion, the availability of data from 244,848 women allowed us to assess the phenotypic penetrance of traits associated with X chromosome aneuploidy in an adult population setting. Our results suggest that the clinical management of women with 45,X/46,XX mosaicism should be minimal, particularly those identified incidentally.

**Funding:** None

## Introduction

X chromosome aneuploidy is a common chromosome abnormality that may be detected as an incidental finding^1^. It can be difficult to determine the clinical importance of the abnormality, as population based studies of X chromosome aneuploidy are usually limited to a small number of cases ascertained from prenatal screening, with follow-up only into young adulthood at the latest^2–4^. Cohorts recruited through identification of clinical features of X chromosome aneuploidy syndromes may suffer from ascertainment bias, because only individuals with those features are being tested. It is increasingly important to determine the phenotypic consequences of genetic abnormalities in non-clinical populations as more genomic testing is carried out on individuals in the general population, including direct to consumer testing, and more extensive whole genome analyses in individuals with recognised conditions^5^.

Women with a 47,XXX chromosome complement are reported to be taller than average and to have earlier menopause, but the evidence to support these associations is limited to potentially biased collections of cases reaching medical attention^6^. Girls with trisomy X are also at increased risk of developmental delay, particularly problems with speech and language^6^. In contrast women with a 45,X chromosome complement or Turner syndrome are generally short (20cm below population mean), may have physical features such as a webbed neck and about 80% have primary amenorrhea^7^. Women with Turner syndrome are at increased risk of hearing difficulties and cardiac disorders, particularly dissection of the aorta^8,9^ ^10,11^. Hypergonadotrophic hypogonadism in Turner syndrome means that pregnancy is often difficult to achieve spontaneously, but women may opt for *in vitro* fertilisation with donor eggs. However, pregnancy in women with Turner syndrome is considered high risk, predominantly due to the increased risk of cardiac complications^12^.

Trisomy X occurs in approximately 1/1,000 newborn females and Turner syndrome in about 1/2,500^13^. Both karyotypes can exist in mosaic form with a mixture of aneuploid and 46,XX cell types in varying proportions. The presence of a normal 46,XX cell line is generally associated with a less severe phenotype, but may depend on tissue-specific variability in degree of mosaicism. X chromosome loss can also occur with normal ageing, with 7.3% of lymphocytes having a 45,X karyotype in women over 65 years old^14^. Traditionally X chromosome aneuploidy was detected by cytogenetic testing, which is expensive and labour intensive, making large population based studies unfeasible. In this study, we tested over 244,000 women from the UK Biobank, a population-based study of adults aged 40-70, for copy number variation using SNP array data and identified 326 individuals with whole X chromosome imbalances. The availability of such a large population sample allowed us to characterise with high statistical confidence the phenotypic characteristics, including adult diseases, of women with X chromosome aneuploidy in a non-clinical context.

## Material and Methods

### UK Biobank cohort

UK Biobank recruited over 500,000 individuals aged 37-73 years (99.5% were between 40 and 69 years) between 2006–2010 from across the UK. Participants provided a range of information via questionnaires and interviews (e.g. demographics, health status, lifestyle) and anthropometric measurements, blood pressure readings, blood, urine and saliva samples were taken for future analysis: this has been described in more detail elsewhere^15^. SNP genotypes were generated from the Affymetrix Axiom UK Biobank array (~450,000 individuals) and the UKBiLEVE array (~50,000 individuals). This dataset underwent extensive central quality control (http://biobank.ctsu.ox.ac.uk). We based our study on 451,099 individuals of white European descent as defined by principal component analysis (PCA). Briefly, principal components were generated in the 1000 Genomes Cohort (phase 3) using high-confidence SNPs to obtain their individual loadings. These loadings were then used to project all of the UK Biobank samples into the same principal component space and individuals were then clustered using principal components 1 to 4. We removed 7 participants who withdrew from the study, and 348 individuals whose self-reported sex did not match their genetic sex based on relative intensities of X and Y chromosome SNP probe intensity. These steps resulted in 244,848 females who were carried forward for subsequent analyses. Basic characteristics for these women are given in **Supplemental Table S1**.

### Identification of women with chromosome X aneuploidy

Log R Ratio (LRR) and B Allele Frequency (BAF) values for 18,725 non-pseudoautosomal, chromosome X SNP probesets were provided by UK Biobank. For each female, we calculated the mean LRR, and the number of probesets falling inside the expected BAF heterozygous range (values between 0.49 and 0.51). We classified females as having X chromosome aneuploidy if they were outliers in both the cohort mean LRR distribution and the expected heterozygous BAF count distribution (**Supplemental Fig. S1**). Women who were outliers for both mean-LRR and BAF heterozygote count were identified using the standard definition of a value more than 1.5 times the interquartile range away from the mean value in the study^16^. We visually inspected the distribution of chromosome-wide LRR and BAF to ensure aneuploidy was consistent along the full X chromosome only (Fig. 1, **Supplemental Figs S2-S4**). Finally, we checked for relatedness amongst the X chromosome aneuploidy cases and we found no UK Biobank defined 1^st^-3^rd^ degree related individuals.

**Figure 1.**
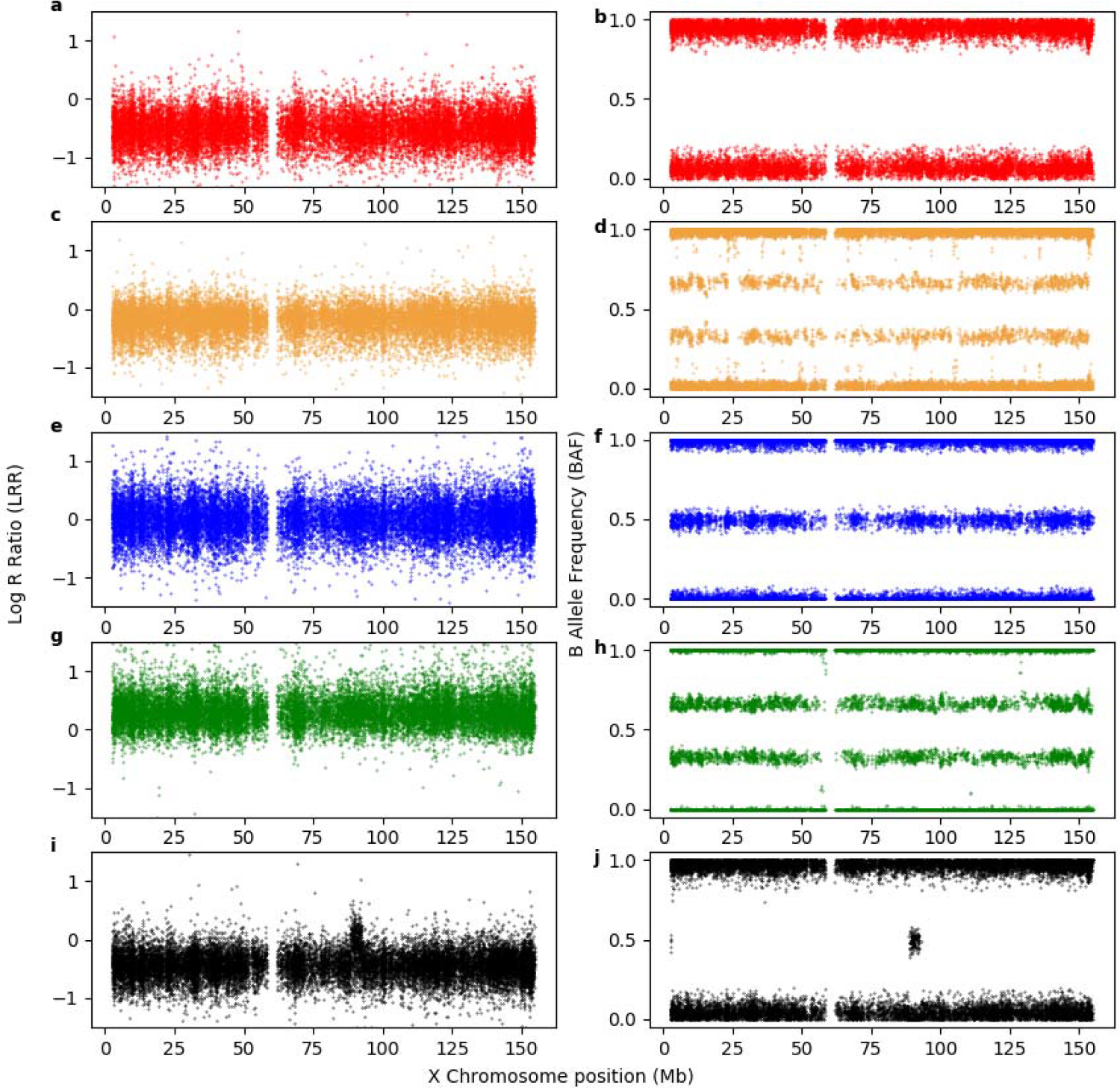
Exemplar Log R Ratio (LRR) and B Allele Frequency (BAF) plots for each detected X chromosome state. (a) and (b) show the respective LRR and BAF for a >80% dosage 45,X individual, (c) and (d) represent the LRR and BAF for a mosaic (≤80%) dosage 45,X/46,XX individual respectively, (e) and (f) represent the respective LRR and BAF of a 46,XX individual and (g) and (h) represent the respective LRR and BAF of a 47,XXX individual. Finally for comparison, plots (i) and (j) represent the respective LRR and BAF of a 46,XY male illustrating a similar LRR dosage to 45,X - note the 4.5Mb 2-copy dosage region at 88.5Mb on chromosome X which is homologous to the Y chromosome present in all males (PAR3).

In females with chromosome X loss, the percentage of mosaicism was estimated by dividing the mean LRR for each woman, by the lowest observed mean LRR across all analysed individuals, giving a ‘dosage’ of 45,X cells. We defined females with chromosome X loss as 45,X/46,XX if the dosage was ≤80% and as 45,X if the dosage was >80%.

We repeated our method of X chromosome aneuploidy identification to detect women with either a partial X chromosome deletion (46,XX,del(X)) or an isochromosome Xq karyotype (46,X,i(Xq)) by analysing probes on Xp and Xq separately, rather than averaging over the whole X chromosome. Those that had a deletion of Xp and duplication of Xq were classified as 46,X,i(Xq) and those with a deletion of either Xp or Xq were classified as 46,XX,del(X).

### Validation of methodology

To validate the use of SNP genotype dosage for determining 45,X mosaicism, we tested six lymphocyte DNA samples from an independent source. The DNA was from individuals tested at the Wessex Regional Genetics Laboratory with a range of 45,X mosaicism, as determined by traditional karyotyping. The samples included a non-mosaic 45,X sample, a 46,XX sample and four 45,X/46,XX mosaics. The six samples were genotyped using both the Illumina Infinium HTS assay on Global Screening array, and the Affymetrix Axiom UK

Biobank array. Processing and derivation of LRR and BAF were carried out using Illumina Genome Studio for the Illumina array data. PennCNV-Affy was used to normalise and derive LRR/BAF values from the raw probe intensity data from the Affymetrix array together with 1,000 randomly selected UK Biobank samples. The percentage dosage was derived for Illumina and Affymetrix data separately by dividing mean LRR by the smallest detected mean LRR value amongst the 6 samples. All analysis was carried out blinded to the results of the karyotyping analysis. We checked for concordance between the independent karyotyping results and mean LRR across chromosome X and then visually inspected the LRR/BAF plots (**Supplemental Fig. S5**).

### Association testing for a range of phenotypes

A range of phenotypes were available in UK Biobank, derived from self-reported questionnaire data, ICD-10 diagnoses recorded in Hospital Episodes Statistics (HES) data and measurements taken at baseline clinic visits as part of the study. ICD-10 diagnoses are added to HES if an individual is admitted to hospital as an inpatient. Seventy-eight percent of UK Biobank individuals had a HES record with one or more ICD-10 codes. We tested the association of X chromosome aneuploidy with a range of phenotypes and traits including: anthropometric, reproductive, cardiovascular, learning/memory and incidence of various diseases (**Supplemental Table S1**). X chromosome aneuploidy was stratified into 4 groups: all 45,X and 45,X/46,XX samples combined, 45,X samples only, 45,X/46,XX samples only and 47,XXX samples. Controls were all other women in UK Biobank (max N=244,522). Association testing was carried out with inverse normalised phenotypes to account for any skewed distributions, using linear regression models in STATA 13, adjusting for SNP chip type (UKB Axiom or UK BiLEVE), ancestry-principal components 1 to 5 supplied by UK Biobank, test centre and age (or year of birth for age at menarche) with the exception of three traits: hypertension, hypothyroidism and household income. Hypertension and hypothyroidism were tested using logistic regression and household income was tested using ordinal logistic regression. The logistic and ordinal logistic models were adjusted using the same covariates as the linear regression model.

## Results

### Detection of chromosome X aneuploidy

We detected 326 women with whole X chromosome imbalances among the 244,848 women we analysed from the UK Biobank, 110 with a higher dosage indicating an additional X chromosome and 216 with a lower dosage, suggesting only one X chromosome in some or all cells (Fig. 1 **and Supplemental Figs S2-S4**). Of the 216 suspected 45,X women there was a range of estimated dosages with mean LRRs ranging from −0.5 to −0.07 (mean = −0.23). Of these women, 30 had an estimated X chromosome dosage in the lowest quintile based on LRR, suggesting the majority of cells had only one X, consistent with a 45,X karyotype. The prevalence of 45,X Turner syndrome detected by SNP array in UK Biobank was therefore 1/8,162 women. Our classification of 186 women with a dosage ≤ 80% as mosaic, gives a prevalence of 1/1,316 women with a mosaic 45,X/46,XX karyotype in UK Biobank.

### Detection of 45,X and 45,X/46,XX mosaicism compared to hospital record data

Hospital record (HES) data was available for 78% of UK Biobank participants at the time of analysis and was the only source of additional Turner syndrome diagnosis information, because no women recorded having Turner syndrome in the self-reported data at their baseline UK Biobank assessment. Of the 30 women we classified with non-mosaic 45,X, 29 had HES data available, of whom 16 had an ICD-10 code for Turner syndrome. This difference left thirteen individuals who had an overnight hospital stay, but for whom Turner syndrome was not recorded (**Supplemental Fig. S6**). It is possible that Turner syndrome was not relevant to the reason for their hospital visit and was therefore not documented. Full details of how the women we detected as non-mosaic 45,X compared to those in the hospital record data are given in **supplemental fig. S6**. Of the 186 women we classified as having mosaic 45,X/46,XX only one had a HES record of ‘Turner syndrome’, and she had an estimated X chromosome loss of 77.5%, very close to our cut off of 80% for calling non-mosaic 45,X Turner syndrome.

In addition to the 216 women we characterised with SNP array data as 45,X or 45,X/46,XX mosaic, 7 women had a HES record of either ‘Turner syndrome’ (n=5) or ‘Mosaicism 45,X/46,XX’ (n=2). Of the five women with a HES record of non-mosaic ‘Turner syndrome’, two were not outliers for whole X chromosome dosage, but on inspection of their SNP array intensity and BAF data, they had a deletion of the p arm and duplication of the q arm, suggesting 46,X,i(Xq) (**Supplemental Fig. S7**); and three were not detected by the SNP array method because their SNP array metrics were well within the normal range (in each case, an above average LRR and an above average frequency of heterozygous BAF). We have no details about the previous diagnosis in these 3 women, e.g. whether diagnosis confirmed by cytogenetics or just based on clinical features. It is possible that some HES codes were entered in error. The two women with a HES record of ‘Mosaicism 45,X/46,XX’ both had a normal 46,XX array profile with LRR and frequency of heterozygous BAF above the 51st centile. These samples were either below the limit of detection or could represent age-related loss of the X which is not detected by our array method.

### Detection of partial X chromosome deletion and isochromosome Xq

After detecting that some women with HES records of ‘Turner syndrome’ had a likely isochromosome Xq, we reanalysed the data separately for the X chromosome p and q arms to detect additional cases with this form of Turner syndrome. We identified a total of five potential 46,X,i(Xq) cases, including the two that had a ‘Turner syndrome’ ICD-10 code (**Supplemental Fig. S8**). This protocol modification also identified nine women with an X chromosome deletion of between 15 and 55 megabases in size (**Supplemental Fig. S9**). The 14 women with partial deletions and 46,X,i(Xq) were of shorter stature (**Supplemental Table S2**).

### Validation of 45,X mosaicism

For the 186 women we classified as mosaic 45,X/46,XX, the estimated percentage of 45,X cells per sample ranged from 15-79%. We validated the accuracy of the SNP array method for determining 45,X mosaicism by testing 6 independent samples which had been tested by conventional cytogenetic techniques, using two different SNP arrays (Table 1, **Supplemental Fig. S5**). There was high level of concordance between the arrays and cytogenetic estimates, with all samples differing by <10%. A perfect correlation between the two methods may be unrealistic as cytogenetic analysis is carried out on cultured lymphocytes, rather than whole blood and thus could be subject to clonal expansion of one cell population over the other. We predicted that age-related loss of the X chromosome would not be detected by the SNP array methodology. As age-related loss is a random process, with a different X being lost in each cell, the BAF pattern would not differ from 46,XX women and the change in dosage as measured by LRR would be relatively small and beyond the limit of detection by our method. However, the SNP array method can detect 45,X mosaics representing a clonal 45,X cell line as the same X chromosome will be absent in each 45,X cell. In summary we were able to verify that the SNP array methodology was able to accurately call 45,X genetic abnormalities down to about 15% 45,X cells, but not age-related loss of the X.

**Table 1.** Degree of X chromosome loss in samples tested by conventional cytogenetics, compared to SNP arrays.

| **Sample** | **Cytogenetic**<br>**estimate** | **Illumina array estimate** | **Affymetrix array**<br>**estimate** |
| --- | --- | --- | --- |
| 1 | 100% 45,X | 100% 45,X (**Fig. S1a**) | 100% 45,X (**Fig. S1b**) |
| 2 | 11% 45,X* | 2% 45,X (**Fig. S1c**) | 13% 45,X (**Fig. S1d**) |
| 3 | 26% 45,X | 25% 45,X (**Fig. S1e**) | 23% 45,X (**Fig. S1f**) |
| 4 | 46,XX | 46,XX (**Fig. S1g**) | 46,XX (**Fig. S1h**) |
| 5 | 24% 45,X | 29% 45,X (**Fig. S1i**) | 30% 45,X (**Fig. S1j**) |
| 6 | 50% 45,X | 48% 45,X (**Fig. S1k**) | 67% 45,X<sup>+</sup> (**Fig. S1l**) |

\* Cytogenetics report stated that this could be due to age-related loss of the X, this was the only control sample from a patient possibly old enough to demonstrate age-related loss (34 years old) and was referred in her mid-thirties for infertility. Both SNP arrays estimated a low level of mosaicism consistent with the cytogenetic estimate, but the LRRs and BAFs were not sufficiently outside the normal range for a sample such as this to be included in our case series. Sample 2 therefore either represents a sample outside the level of detection for this method, or that this individual has age-related loss of the X <sup>14</sup>.

### 45,X and 45,X/46,XX mosaicism and height

The 30 women with evidence of non-mosaic 45,X chromosome loss were on average 17.2cm shorter than 46,XX females (Tables 2 and 3, Fig. 2, **Supplemental Table S3**). In contrast, the 186 mosaic 45,X/46,XX cases were on average only 3.8cm shorter than 46,XX women (*P* = 4.0 x 10^−14^). Height ranged from 139-182cm in the mosaic group, and 7% of these women would be taller than one standard deviation above the mean height in 46,XX women. There was a weak correlation between percentage mosaicism and height in the women with mosaic 45,X/46,XX (r=0.23).

**Table 2.**
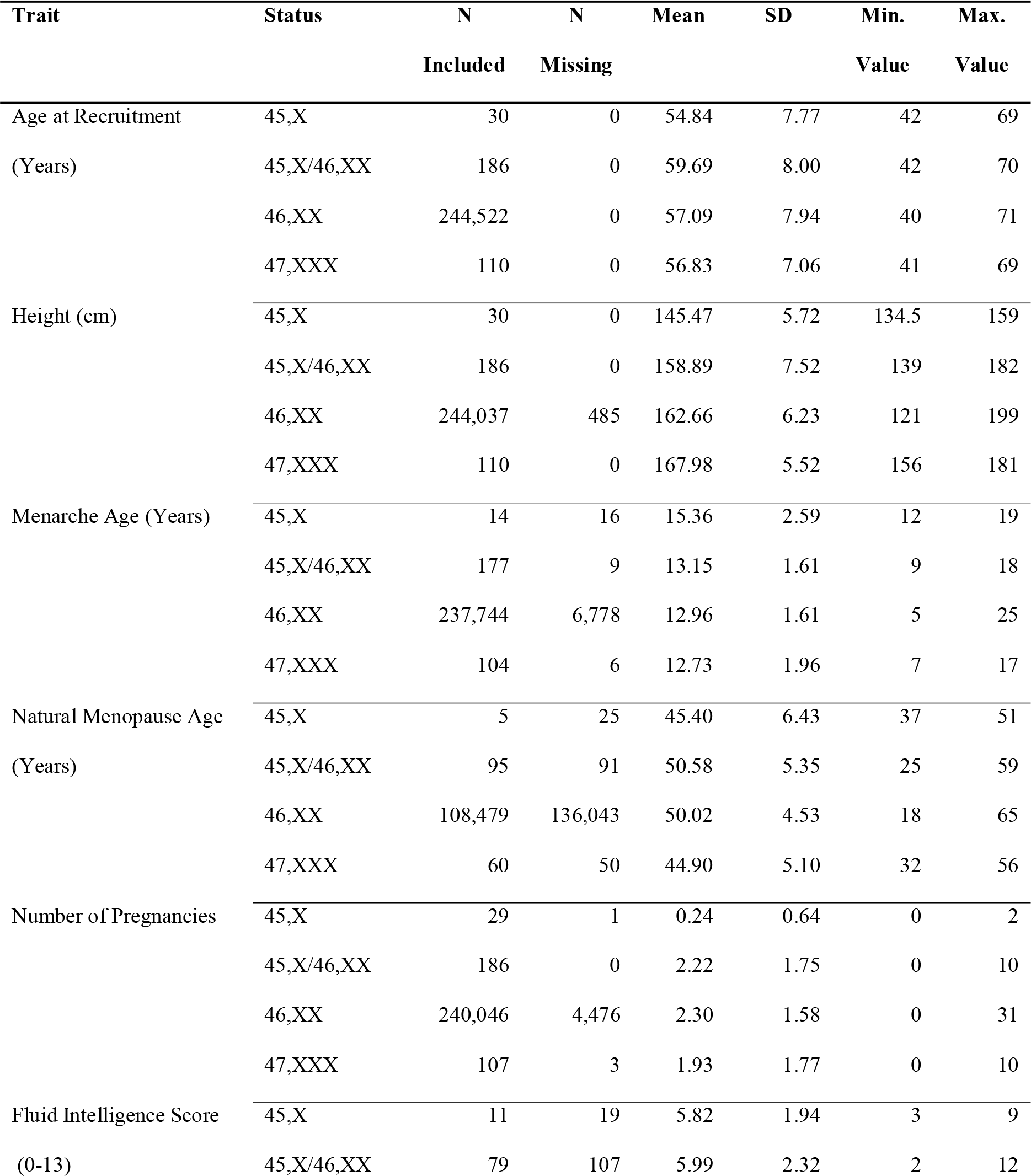
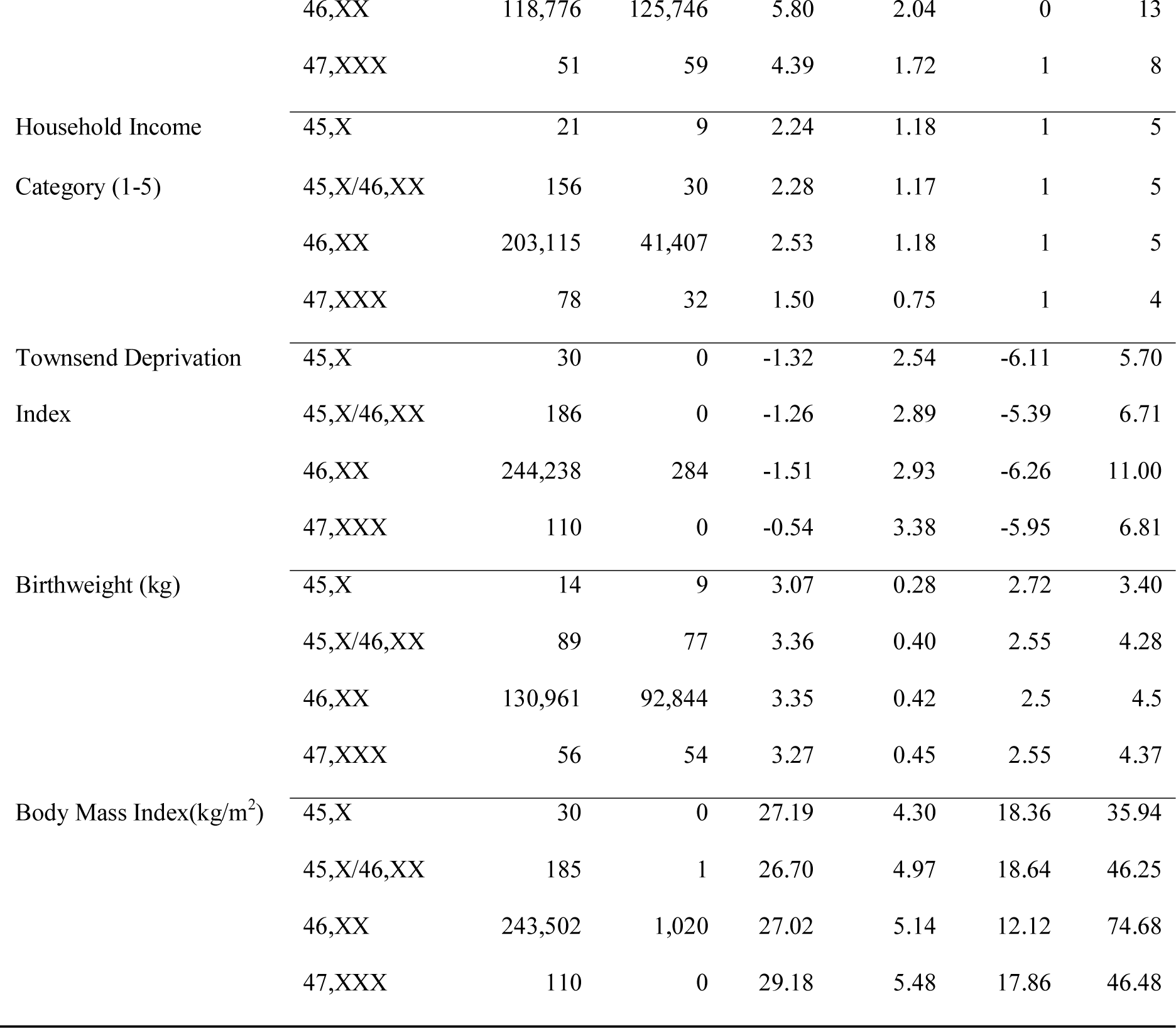
Characteristics of UK Biobank females stratified by X chromosome aneuploidy status. ‘45,X’ are individuals assumed to be non-mosaic Turner syndrome individuals. ‘45,X/46,XX’ are individuals assumed to be mosaic Turner syndrome. ‘46,XX’ are controls, and ‘47,XXX’ are trisomy X individuals. 16 and 25 non-mosaic 45,X cases did not report menarche or menopause ages respectively. (N=number of individuals included in analysis, SD = standard deviation)

**Table 3.** Association between X chromosome aneuploidy and nine phenotypes. The effect of the aneuploidy on each trait is given in number of standard deviations change compared to 46,XX women, with an association *P* value (N=number of cases included in analysis, SD = standard deviation, CI = confidence interval). *P* values that pass the threshold for statistical confidence, corrected for multiple testing, are highlighted (i.e. Equivalent to *P*<0.05).

| Trait | Status | N | Effect<br>(SD) | Low 95%<br>CI (SD) | High 95%<br>CI (SD) | *P* |
| --- | --- | --- | --- | --- | --- | --- |
| Height (cm) | 45,X | 30 | -2.73 | -3.07 | -2.38 | **$1.5 \times 10^{-53}$** |
|  | 45,X/46,XX | 186 | -0.54 | -0.68 | -0.40 | **$4.0 \times 10^{-14}$** |
|  | 47,XXX | 110 | 0.84 | 0.66 | 1.03 | **$5.8 \times 10^{-20}$** |
| Menarche Age (Years) | 45,X | 14 | 1.33 | 0.81 | 1.86 | **$5.9 \times 10^{-7}$** |
|  | 45,X/46,XX | 177 | 0.12 | -0.03 | 0.26 | 0.12 |
|  | 47,XXX | 104 | -0.15 | -0.34 | 0.05 | 0.14 |
| Natural Menopause Age<br>(Years) | 45,X | 5 | -0.99 | -1.86 | -0.13 | 0.024 |
|  | 45,X/46,XX | 95 | 0.12 | -0.08 | 0.32 | 0.23 |
|  | 47,XXX | 60 | -0.98 | -1.23 | -0.73 | **$1.2 \times 10^{-14}$** |
| Number of<br>Pregnancies | 45,X | 29 | -1.50 | -1.86 | -1.14 | **$3.3 \times 10^{-16}$** |
|  | 45,X/46,XX | 186 | -0.12 | -0.27 | 0.02 | 0.09 |
|  | 47,XXX | 107 | -0.26 | -0.44 | -0.07 | 0.0076 |
| Fluid Intelligence Score<br>(0-13) | 45,X | 11 | -0.17 | -0.74 | 0.39 | 0.55 |
|  | 45,X/46,XX | 79 | 0.09 | -0.12 | 0.30 | 0.39 |
|  | 47,XXX | 51 | -0.74 | -1.00 | -0.48 | **$3.7 \times 10^{-8}$** |
| Household Income<br>Category (1-5) | 45,X | 21 | -0.89 | -1.66 | -0.12 | 0.024 |
|  | 45,X/46,XX | 156 | -0.21 | -0.51 | 0.08 | 0.15 |
|  | 47,XXX | 78 | -1.98 | -2.45 | -1.51 | **$8.7 \times 10^{-17}$** |
| Townsend Deprivation<br>Index | 45,X | 30 | 0.13 | -0.20 | 0.47 | 0.43 |
|  | 45,X/46,XX | 186 | 0.13 | 0.00 | 0.27 | 0.052 |
|  | 47,XXX | 110 | 0.24 | 0.07 | 0.42 | 0.0067 |
| Birthweight (kg) | 45,X | 13 | -0.67 | -1.19 | -0.14 | 0.013 |
|  | 45,X/46,XX | 89 | 0.02 | -0.19 | 0.23 | 0.86 |
|  | 47,XXX | 56 | -0.21 | -0.47 | 0.05 | 0.11 |
| Body Mass Index (kgm<sup>2</sup>) | 45,X | 30 | 0.13 | -0.26 | 0.51 | 0.51 |
|  | 45,X/46,XX | 185 | -0.09 | -0.25 | 0.06 | 0.23 |
|  | 47,XXX | 110 | 0.46 | 0.26 | 0.66 | **6.3 x 10<sup>-6</sup>** |

**Figure 2a.**
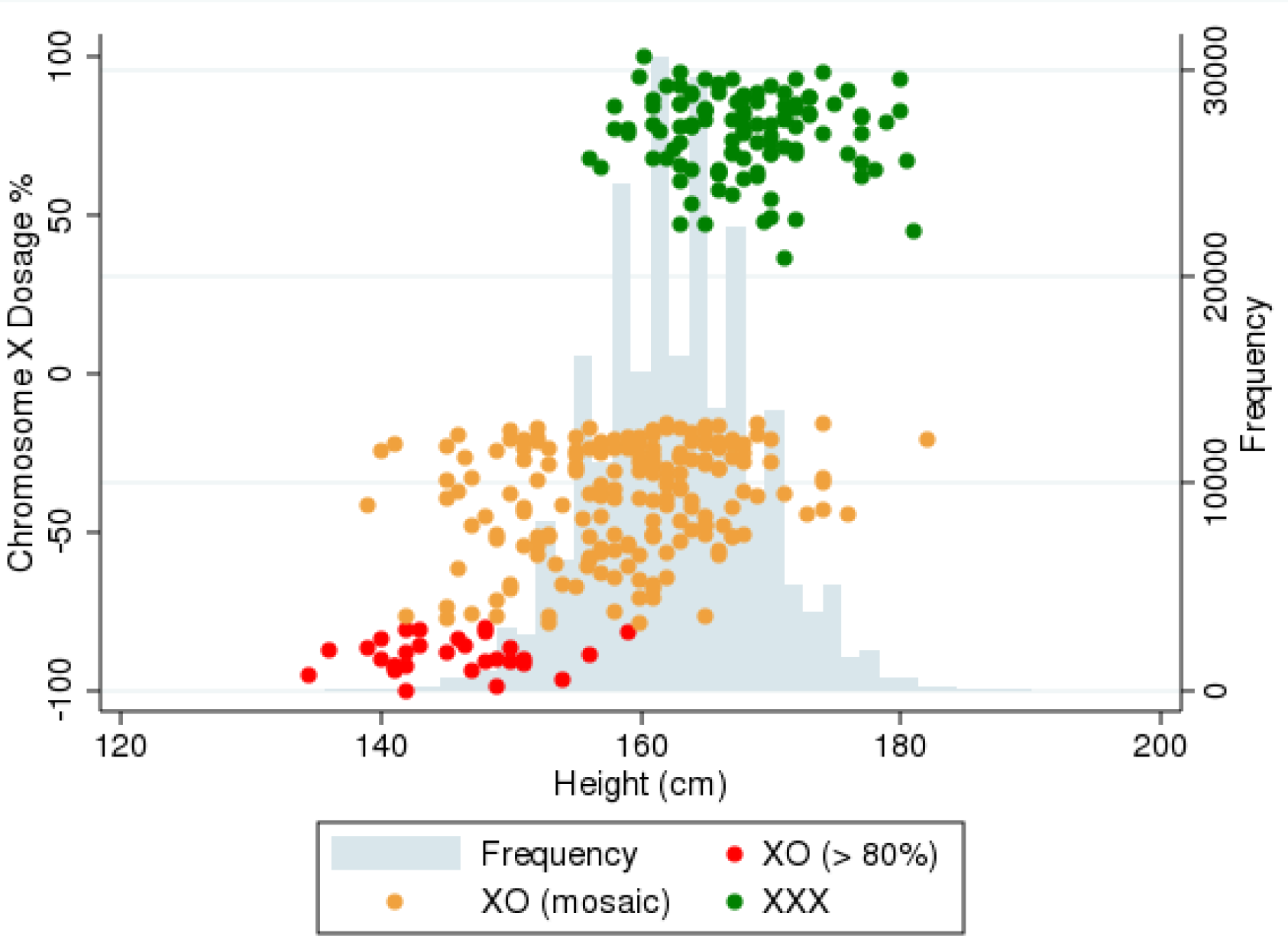
Relationship between chromosome X dosage and height in 45,X and 47,XXX females. X chromosome dosage is colour coded (see key) for each category tested in the analysis. Distribution of height in controls is shown in grey histogram.

**Figure 2b.**
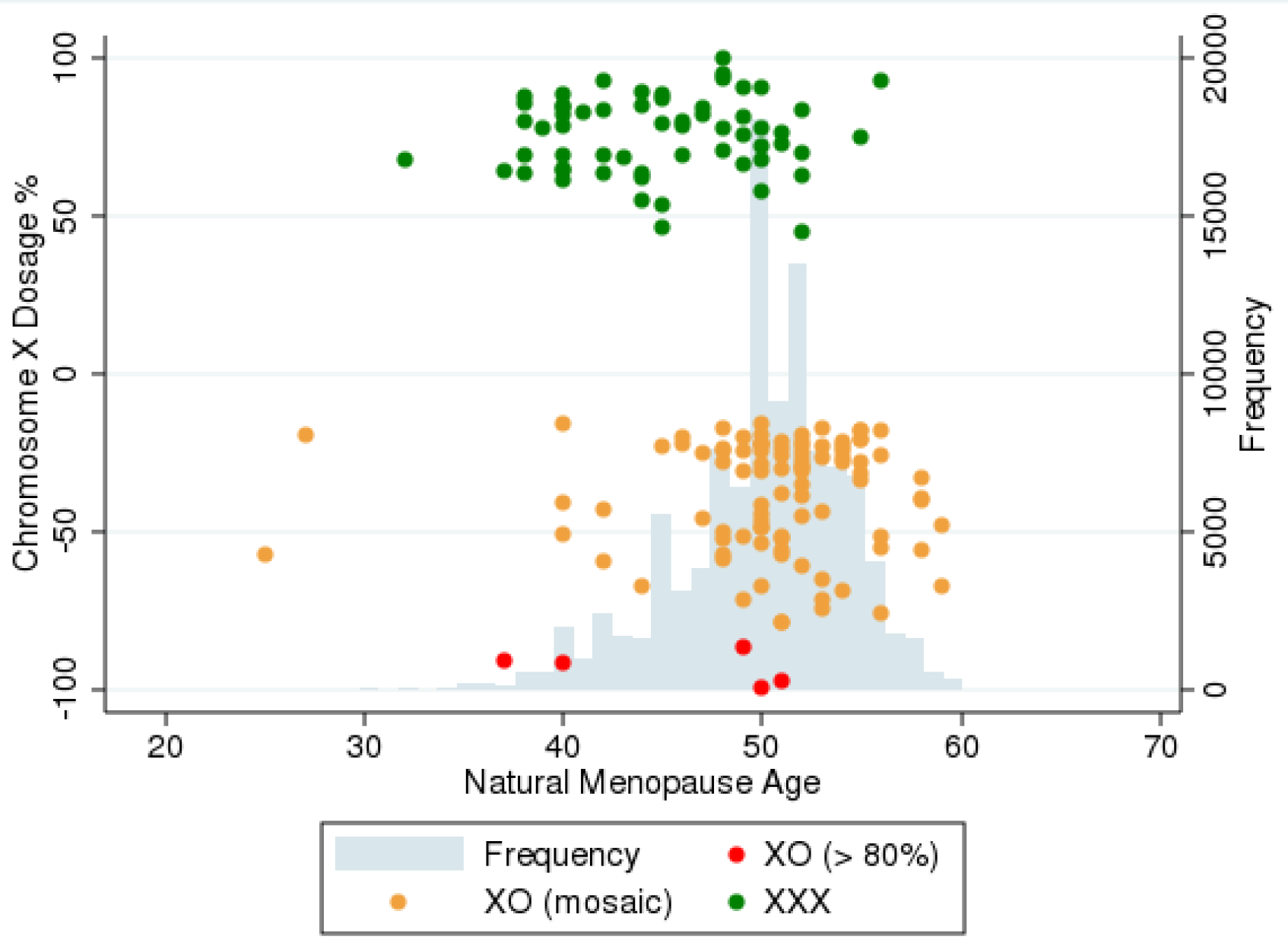
Relationship between chromosome X dosage and natural menopause age in 45,X and 47,XXX females. X chromosome dosage is colour coded (see key) for each category tested in the analysis. Distribution of menopause age in controls is shown in grey histogram.

### 45,X and 45,X/46,XX mosaicism and ovarian function

Primary amenorrhea was not specifically coded in UK Biobank, but 16 of the 30 women with non-mosaic 45,X answered ‘don’t know’ or ‘Prefer not to answer’ to the question “How old were you when your periods started?”, which were the only options apart from an age in years. This proportion (53%) was much higher than that in 46,XX women (2.9%) with an odds ratio for selecting “don’t know” or “prefer not to answer” compared to 46,XX women of 39.5 (95% CI = 18.1, 87.7; *P* = 1.5 x 10^−17^). We therefore assumed that a large proportion of these women did not go through menarche. For the other 14 women with non-mosaic 45,X, menarche ages of between 12 and 19 years were recorded. It is possible that menarche was induced with exogenous hormones in these 14 individuals, but there was no data to indicate this. In contrast, only 9 of the 186 women with mosaic 45,X/46,XX did not report an age at menarche, and the mean menarche timing was 13.2 years in those 177 individuals who recorded an age, not different from the mean age of 12.95 years in the 46,XX women (*P* = 0.12).

The recruitment ages of the 30 women with non-mosaic 45,X ranged from 42-69 years, but only 5 reported a natural menopause age and the odds ratio for not reporting an age at natural menopause compared to women with 46,XX was 3.98 (95% CI = 1.50, 13.31; *P* = 0.0026). Ninety-five of the 45,X/46,XX mosaic women reported an age at natural menopause (Tables 2 and 3, Fig. 2, **Supplemental Table S3**), at an average age of 50.6 years (range 25-59 years), again not different from the mean age in control women with 46,XX (mean = 50 years (range 18-65); *P* = 0.23).

Only four of the thirty non-mosaic 45,X cases had ever been pregnant, much fewer than in the remaining 46,XX women with an odds ratio for never being pregnant of 32.35 (95% CI = 11.21, 127.22; *P* = 8.98 x 10^−17^). Most of the mosaic 45,X/46,XX cases reported a pregnancy, with only 37 of 186 reporting they had never been pregnant, a frequency not different from women with 46,XX. The mean number of pregnancies in the 45,X/46,XX women was 2.2, not different from the control 46,XX individuals (*P* = 0.09). There was also no increased incidence of pregnancy loss in the women with 45,X/46,XX, either from miscarriage, termination or stillbirth.

### 45,X and 45,X/46,XX mosaicism and cardiac function

Heart defects are a common reported feature of Turner syndrome, but we found no substantially increased incidence of heart defects in this cohort of 45,X women, based on self-reported incidence of disease and ICD-10 codes (e.g. heart valve disease OR = 8.08; 95% CI = 0.93, 32.08; *P* = 0.005). Blood pressure and hypertension were elevated (e.g. hypertension OR = 3.15; 95% CI = 1.41, 6.99; *P* = 5 x 10^−3^), and arterial elasticity was 1.05 indices lower in the 30 non-mosaic 45,X women (se = 0.32; *P* = 0.001). Four of the 30 women with 45,X had a medically diagnosed heart condition, including coarctation/dissection of the aorta and myocardial infarction and 13 were on blood pressure medication (OR = 3.66; 95% CI = 0.93, 32.08; *P* = 7.7 x 10^−4^). Hypertension and blood pressure were not raised in the mosaic 45,X/46,XX women (hypertension OR = 1.02; 95% CI = 0.75, 1.40; *P* = 0.88) (Table 4), The women with mosaic 45,X/46,XX had not had more cardiac operations and were not more likely to be on blood pressure medication than the 46,XX women.

**Table 4.**
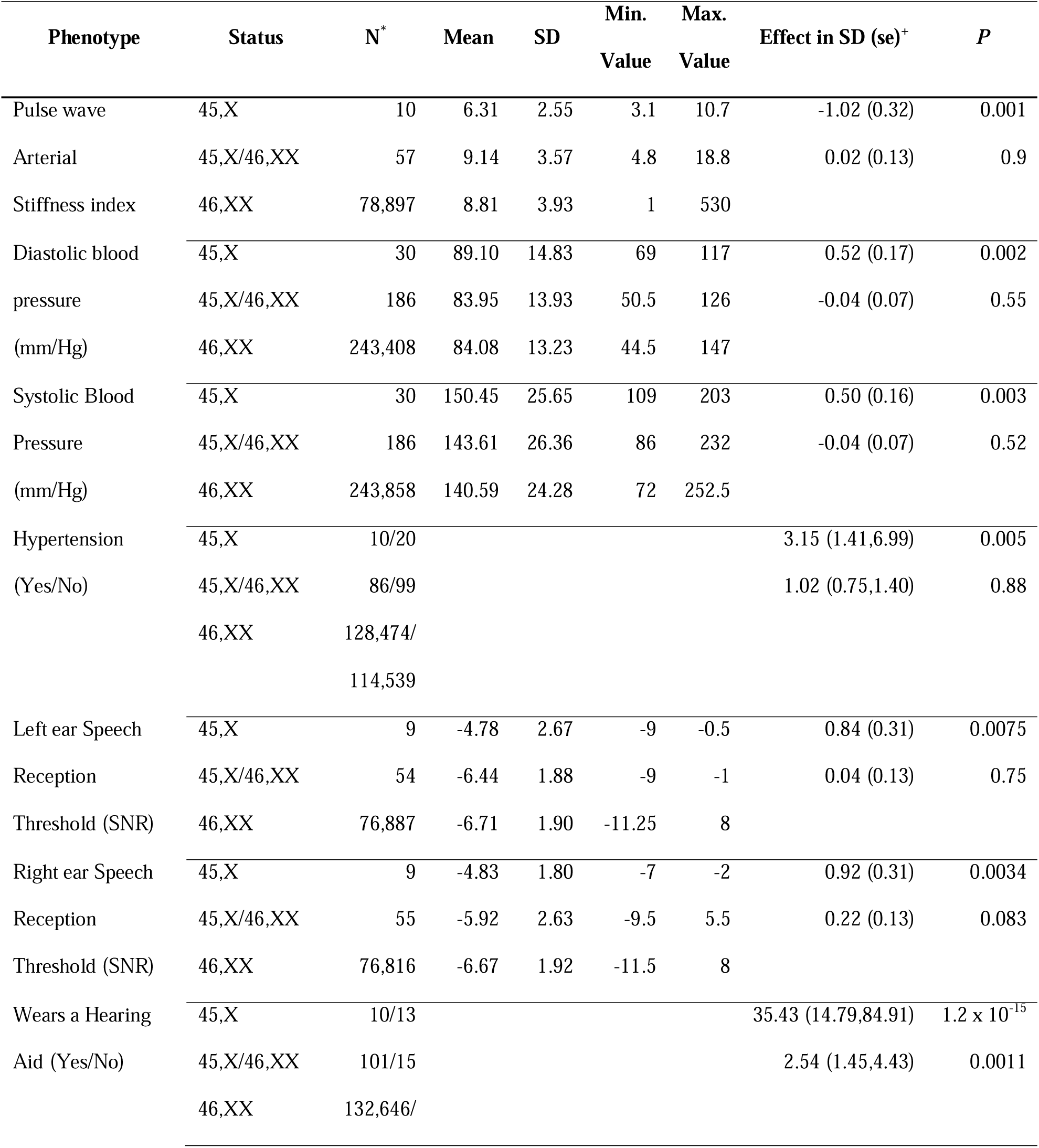
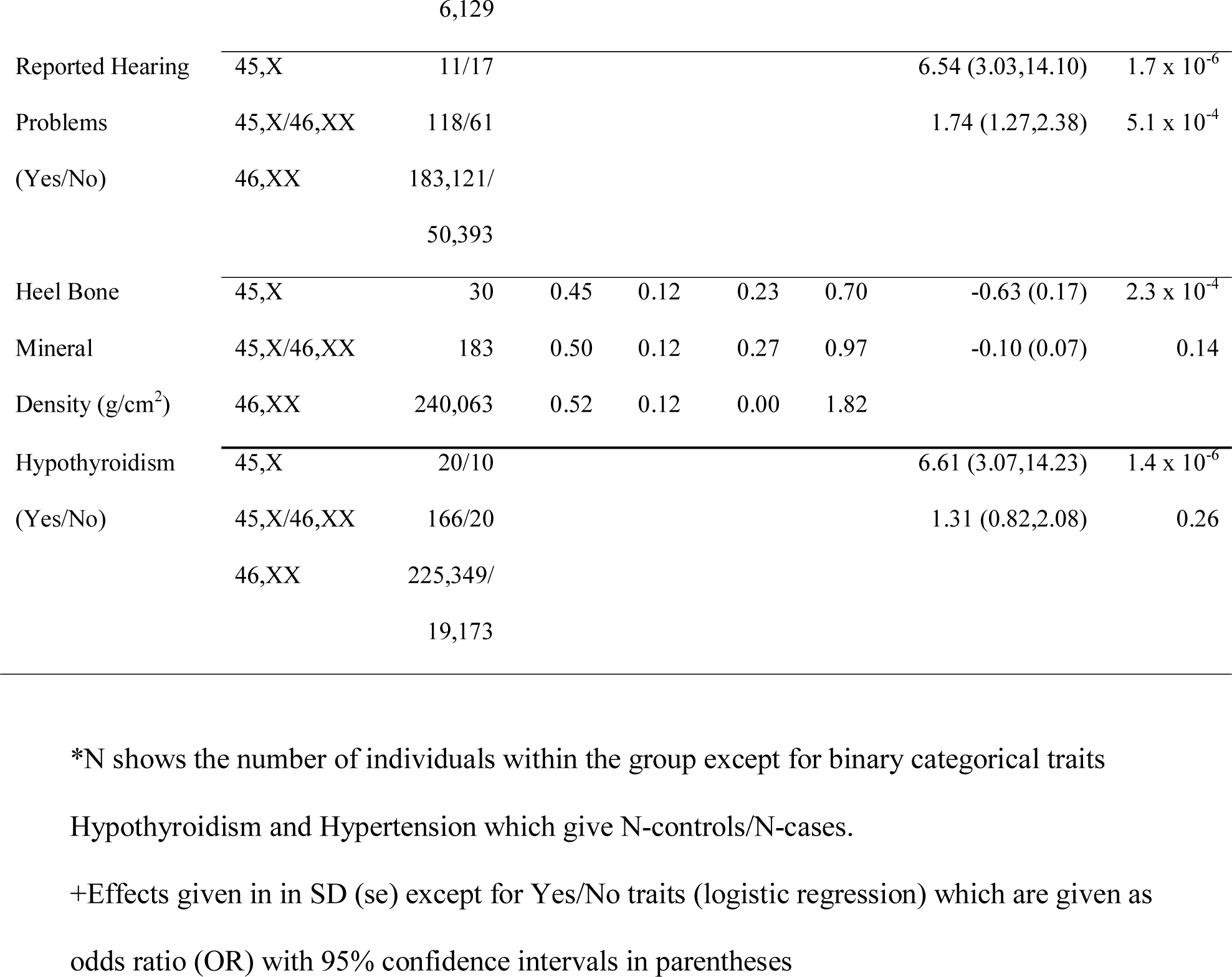
Arterial stiffness, blood pressure, deafness, heel bone mineral density, hypothyroidism and hypertension in women with X chromosome aneuploidy compared to women with 46,XX karyotype. The effect of the aneuploidy on each trait is given in standard deviations compared to 46,XX women with an association *P* value (N = number of individuals included in analysis, SD = standard deviation, se = standard error).

### Additional disorders in women with X chromosome aneuploidy

We detected several other conditions in the women with non-mosaic 45,X, but few in the women with mosaic 45,X/46,XX. There was an increased prevalence of diagnosed hypothyroidism in the non-mosaic 45,X women compared with 46, XX women (OR = 6.61; 95% CI = 3.07, 14.23; *P* = 1.4 x 10^−6^). Heel bone mineral density was slightly reduced in the non-mosaic 45,X women compared to 46,XX women (ß = −0.63 SDs; se = 0.17 SDs; *P* = 2.3 x 10^−4^). There was an increased incidence of hearing abnormalities in the non-mosaic 45,X women, with more women wearing a hearing aid (OR = 35.43; 95% CI = 14.79, 84.91; *P* = 1.2 x 10^−15^), reporting ‘hearing problems’ (OR = 6.54; 95% CI = 3.03, 14.10; *P* = 1.7 x 10^−6^) and having poorer average measured hearing than 46,XX women (ß = 0.88 SDs; se = 0.31 SDs; *P* = 0.0055) (Table 4). In contrast there was no increased prevalence of diagnosed hypothyroidism in the mosaic 45,X/46,XX women (OR = 1.31; 95% CI = 0.82, 2.08; *P* = 0.26) and bone mineral density was similar to that of 46,XX women (Table 4). We did observe hearing problems in the mosaic 45,X/46,XX women, but with lower incidence and effect sizes. For example, there were more 45,X/46,XX women wearing a hearing aid (OR = 2.54; 95% CI = 1.45, 4.43; *P* = 1.1 x 10^−3^) or reporting ‘hearing problems’ (OR = 1.74; 95% CI = 1.27, 2.38; *P* = 5.1 x 10^−4^) than 46,XX women, but no difference in measured hearing (*P* > 0.05) (Table 4).

### Trisomy X phenotype

Trisomy X was detected in 110 women, giving a prevalence of 1/2,226 in UK Biobank women. A further two cases appeared to be mosaic 46,XX/47,XXX, but were excluded from further analyses because there were too few in this category to analyse separately. Of the 110 47,XXX women, 90 were present in the HES records, but none had a prior diagnosis of ‘Karyotype 47,XXX’ as determined by ICD-10 code.

We found an association between trisomy X and adult height (ß = 0.84 SDs; 95% CI = 0.66, 1.03; *P* = 5.8 x 10^−20^). Women with 47,XXX were 5.3cm taller on average than 46,XX women (Fig. 2, Tables 2 and 3, **Supplemental Table S3**). Women with trisomy X had normal age at menarche, but an average 5.12 years earlier age of natural menopause (ß = - 0.98 SDs; 95% CI = −1.23, −0.73; *P* = 1.2 x 10^−14^) (Tables 2 and 3, Fig. 2, **Supplemental Table S3**). Women with 47,XXX had a similar number of pregnancies (mean = 1.9, range 0-10) and no higher frequency of pregnancy loss than 46,XX controls. We also found that women with trisomy X had a lower fluid intelligence (ß = −0.74 SDs; 95% CI = −1.0, −0.48; *P* = 3.7 x 10^−8^) and a decreased household income (ß = −1.98 SDs; 95% CI = −2.45, −1.51; *P* = 8.7 x 10^−17^) than 46,XX women (Tables 2 and 3, **Supplemental Table S3**). Sixty three percent of women in the 47,XXX group had a household income less than £18,000 per year, compared to 24% of 46, XX control women. Women with 47,XXX were more likely to live alone (OR = 2.07; 95% CI = 1.36, 3.12; *P* = 7 x 10^−4^) with 35.2% (37/110) living alone compared to 19.8% of 46,XX women. The women with 47,XXX also had a higher BMI than the 46,XX women, by on average over 2 BMI units (ß = 0.46 SDs; 95% CI = 0.26, 0.66; *P* = 6.3 x 10^−6^).

## Discussion

The availability of more than 244,000 adult women with SNP array data in a population based study enabled us to assess X chromosome aneuploidy phenotypes without the potential ascertainment biases of clinical presentation and with sufficient statistical power to quantify phenotypic effects, including adult onset diseases. In contrast previous studies have either concentrated on clinically ascertained women, or prenatally ascertained cases numbering in the tens and not followed up into adulthood. We identified 326 women in UK Biobank with whole X chromosome dosage anomalies: 216 consistent with having a 45,X or 45,X/46,XX mosaic karyotype and 110 with 47,XXX. We detected only two cases of 46,XX/47,XXX mosaicism, but these are less common than 45,X/46,XX mosaics and would be more difficult to detect with the array method^17^.

### Prevalence of X chromosome aneuploidy

We detected a 45,X or 45,X/46,XX mosaic karyotype in 1/1,134 women in UK Biobank. The majority of these women were mosaic. The published prevalence of Turner syndrome is 1/2,500, however only approximately 60% of those Turner syndrome cases are caused by 45,X or 45,X/46,XX mosaicism, the rest being due to other abnormalities such as deletions or isoXq ^13,18^, thus the prevalence of aneuploidy causing Turner syndrome is around 1/4,200.

This is approximately four times lower than the combined prevalence of 45,X and 45,X/46,XX mosaicism we detected in UK Biobank, suggesting that many cases go undiagnosed. The higher prevalence in UK Biobank is explained by the high proportion of mosaic cases in our series, who may not be included in published studies because they have fewer phenotypic features of Turner syndrome and would likely go undetected in clinical practice. Most series report approximately 16% of aneuploidy cases being mosaic 45,X/46,XX, but in UK Biobank the proportion we detected was 86%. The prevalence of 45X was only 1/8,161, lower than the expected prevalence of approximately 1/5,000. This difference is likely to be due in part to the UK Biobank favouring healthy individuals, but also we may have classified some 45,X cases as mosaic. It is also important to note that the mosaic cases we detected are not due to age related X chromosome loss, because this would have resulted in a random loss of paternal or maternal X and a different BAF pattern compared to losing either the maternal or paternal X at, or soon after, conception.

The prevalence of 47,XXX detected by SNP array in UK Biobank was 1/2,226, substantially lower than the reported incidence of 1/1,000 livebirths. This is likely to be due to the UK Biobank favouring healthy individuals^19^. We observed an association with IQ-related measures in women with 47,XXX, and it is known that individuals with lower IQ were less likely to participate in the study^19^.

### Most women with 45,X/46,XX mosaicism detected incidentally will not require clinical follow up

The most important implication of our data is that many women with a 45,X cell line likely do not require currently recommended interventions^20^. The international clinical practice guidelines do not readily differentiate mosaic from non-mosaic 45,X, yet our data suggest that level of mosaicism is an important clinical indicator of Turner syndrome features and related complications. There is less information in the literature about women with a mosaic 45,X/46,XX karyotype, as these generally constitute a smaller proportion of any reported series of cases, and are commonly combined with 45,X cases. In the UK Biobank there were 186 women who appeared to be mosaic for X chromosome loss, i.e. retaining two X chromosomes in a proportion of cells. Surprisingly, the phenotype in these women was unremarkable: while they were slightly shorter on average, many of the women in this category were of average height, with the tallest individual being 182cm, despite 20% of blood cells being 45,X (**Supplemental Fig. S3ah**). This group of women went through menarche and menopause at an average age, had an average number of children, were not at increased risk of pregnancy loss and there was no evidence of increased risk of cardiac complications.

As more individuals are obtaining genomic information either as part of healthcare assessments or through direct-to-consumer genetic testing, the likelihood of detecting X chromosome imbalance is increasing and it is important to be able to counsel individuals appropriately regarding the implications of such findings. Currently any women identified as having a 45,X cell line not thought to be due to age-related loss, would be diagnosed as having Turner syndrome and would be offered extensive monitoring, particularly during pregnancy^20^. While it is recognised that mosaic cases may be less likely to have abnormalities than non-mosaic 45,X Turner syndrome cases they are still considered high risk with respect to cardiac anomalies and hypertension during pregnancy for instance. In young women, one of the main concerns will be the likelihood of infertility associated with a diagnosis of Turner syndrome. Our data suggest that in fact the presence of a 45,X cell line in blood does not adversely affect reproductive lifespan or fertility in most cases, as long as more than 20% of cells have two X chromosomes.

### Of the 326 women in UK Biobank with X chromosome aneuploidy, 30 had a complete loss of one X chromosome, and their features were consistent with a diagnosis of Turner syndrome

Sixteen of the 29 non-mosaic 45,X women with inpatient hospital records had a pre-existing medical diagnosis of Turner syndrome in these records. Only one woman with a previous diagnosis of Turner syndrome was in the mosaic 45,X/46,XX group and this individual was an estimated 77.5% 45,X, confirming that a high proportion of X chromosome loss is required to generate a substantial effect on phenotype, such that would be identified by clinicians. Women with 45,X had a phenotype that was characteristic of that reported for this chromosomal abnormality, i.e. short stature and primary amenorrhea being the main features. Cardiac abnormalities were not very common in this group, but were found in some individuals, confirming the need for continued screening programmes. There were no data available on physical characteristics, such as neck webbing.

### Women with 47,XXX are taller, have earlier menopause and are of below average cognitive ability

These findings are important because they suggest that ascertainment bias has not dramatically influenced the current consensus regarding the 47,XXX phenotype. Our findings, in a cohort without the ascertainment bias of having a clinical referral, agree with published literature on the phenotype associated with 47,XXX syndrome. Traditionally 47,XXX syndrome has been identified by cytogenetic testing, which is a costly and labour intensive procedure and requires fresh tissue to enable culture of dividing cells. Thus the majority of studies on 47,XXX have been on women tested for a clinical presentation such as learning difficulties, or identified through prenatal testing as an incidental finding. Cohorts of older women with 47,XXX are uncommon as prenatal chromosome testing has been relatively recently adopted and thus most longitudinal studies of 47,XXX women ascertained at birth have not yet reached post-menopausal ages. The first reported case of 47,XXX was an infertile woman and infertility or premature ovarian failure is often a reason for referral for 47,XXX testing^21^. However primary ovarian insufficiency (POI) affects 1% of women and 47,XXX is a relatively common finding too (1/1,000) and thus the association could be due to ascertainment bias. Our data suggests that there is a genuine effect on reproductive lifespan in women with 47,XXX. This effect appears to be limited to the end of reproductive life however, as these women had average menarche age, but went through menopause on average more than 5 years earlier than women with 46,XX, with 10 women meeting the criteria for POI, with menopause before 40 years. The frequency of POI in 47,XXX women is therefore 9%, approximately 10 times that seen in 46,XX women. We found no evidence for an impact of 47,XXX on reproductive function throughout life, as these women had an average number of pregnancies and no significant increase in pregnancy loss.

Dosage of the sex chromosomes is known to influence human height^22^. Individuals with a single X chromosome and Turner syndrome are typically on the 0.1 centile for height and an additional X in men with Klinefelter syndrome increases adult height by about one standard deviation. The UK Biobank women with 47,XXX were on average 5.3cm taller than the 46,XX women. Adult height is associated with puberty timing, with earlier puberty resulting in shorter adult height. The 47,XXX effect on adult height is however unlikely to be driven by later puberty timing, as menarche age was not significantly different from the 46,XX women. It is more likely that the increase in height is caused by dosage sensitive genes responsible for post-pubertal growth.

The IQ of girls with 47,XXX is reported to be within the normal range, but approximately 10- 15 points below their sibling average. Lower IQ is also reported in adult cases of 47,XXX, along with psychosocial features, such as low self-esteem, language difficulties and increased prevalence of psychiatric disorders^23^. Few series of older women with 47,XXX have been reported previously, but a study from Denmark found reduced socio-economic status in a series of 108 women with 47,XXX^24^. In the UK Biobank study we found a substantial reduction in fluid intelligence compared to the 46,XX women. ‘Fluid intelligence’ describes the capacity to solve problems that require logic and reasoning ability, independent of acquired knowledge. We observed an increase in BMI in women with 47,XXX, that is not reported as a typical feature of this chromosome abnormality, and may be related to the observed lower IQ. There was also a reduction in household income in women with 47,XXX. The association with lower household income was not a consequence of general lower socio economic status as Townsend deprivation index was not significantly associated with 47,XXX status. Ability to work and type of employment taken by younger women with 47,XXX has been reported in other studies, but with much smaller sample sizes. Thus the reduced IQ observed in adolescents with 47,XXX continues into older age and appear to have a lifelong effect on income.

#### Limitations

The samples tested in this study were from peripheral blood and levels of mosaicism are likely to differ between tissue types. Thus the mosaicism we detected may not reflect the level present in brain, ovary or heart for example. Non-aneuploid causes of Turner syndrome were not included in our analyses, although we detected cases of apparent iso Xq, and X chromosome deletions, but we were unable to verify these cytogenetically. Any cases with Y chromosome material were excluded as sex mismatches.

Our study is also biased in favour of healthier individuals with a more benign phenotype^19^. We know for instance that there are only sixteen cases of Down syndrome in UK Biobank, while we would expect nearer 280 individuals in a population of 500,000 people between 40-70 years old^25^. UK Biobank participants are all over 40 years old and thus cases of Down syndrome may be underrepresented due to reduced life expectancy, nevertheless it is likely that individuals with Down syndrome were less likely to volunteer for the study due to the learning difficulties associated with the condition. Biobank volunteers are more likely to be female, have higher IQ and be from a higher socioeconomic group than the general population^19^. Cases of X chromosome aneuploidy with a more severe phenotype may therefore be underrepresented in the Biobank cohort. Thus our study does not represent a comprehensive assessment of X chromosome aneuploidy in the population, but it goes a long way to redressing the bias seen in many published series of cases ascertained through clinical referrals and therefore more likely to have a clinical phenotype.

#### Validation

While we were not able to karyotype any of the samples from UK Biobank to validate our methodology, we did use two platforms of array-based technology to test an independent series of patients with various levels of 45,X mosaicism and found very close correlation between the array method and cytogenetic assessment. Further validation came from finding 16 previously known Turner syndrome cases in UK Biobank among the 30 individuals who we classified as non-mosaic 45,X with the array dosage data. The array method and analysis pipeline we used will not detect random age-related X chromosome loss and thus the cases we have identified are true clonal mosaics. Karyotype analysis cannot distinguish between age-related loss and presence of a 45,X cell line in patients over 30 years old. Three previously diagnosed cases of Turner syndrome appeared to have a normal 46,XX SNP array profile, this may be a sample mix-up, or a suspected Turner syndrome case that was not confirmed by genetic testing, or perhaps individuals with a more complex genetic abnormality that was not detected by the array method. Indeed two additional previously known Turner syndrome cases were not detected by the SNP array method because they had loss of Xp and gain of Xq making the overall X chromosome dosage not significantly different from normal 46,XX. We suspect that these are isochromosomes, which contribute to about 15% of Turner syndrome cases in other published series^18,26^. We adapted our protocol to find additional cases with non-aneuploid X chromosome abnormalities and found a total of 5 cases of suspected 46X, i(Xq) and 9 with X chromosome deletions, although we were not able to validate these findings with an independent technology. We are confident that the cases we have identified do not include false positives, but there may be X chromosome imbalances with Turner or 47,XXX phenotypes that are not included in our cases.

#### Conclusions

We have assessed X chromosome aneuploidy phenotypes without the potential ascertainment biases of clinical presentation and with sufficient statistical power to quantify phenotypic effects, including adult onset disease. Our study contrasts with previous studies that have either concentrated on clinically ascertained women, or prenatally ascertained cases numbering in the tens and not followed up into adulthood. The main implication of our data is that X chromosome aneuploidy is not always associated with an adverse phenotype: 45,X/46,XX mosaic females had a normal reproductive lifespan and birth rate, with no reported cardiovascular complications compared to 46,XX women. This study presents implications for future management of women with a 45,X/46,XX mosaic karyotype, particularly those identified incidentally.

## Acknowledgements

This research has been conducted using the UK Biobank Resource. This work was carried out under UK Biobank project number 871.

## Ethics UK Biobank

This study was conducted using the UK Biobank resource. Details of patient and public involvement in the UK Biobank are available online (www.ukbiobank.ac.uk/about-biobank-uk/ and https://www.ukbiobank.ac.uk/wp-content/uploads/2011/07/Summary-EGF-consultation.pdf?phpMyAdmin=trmKQlYdjjnQIgJ%2CfAzikMhEnx6). No patients were specifically involved in setting the research question or the outcome measures, nor were they involved in developing plans for recruitment, design, or implementation of this study. No patients were asked to advise on interpretation or writing up of results. There are no specific plans to disseminate the results of the research to study participants, but the UK Biobank disseminates key findings from projects on its website.

## References

1. Otter M, Schrander-Stumpel CT, Curfs LM. Triple X syndrome: a review of the literature. European journal of human genetics: EJHG 2010; 18(3): 265–71.

2. Tokita MJ, Sybert VP. Postnatal outcomes of prenatally diagnosed 45,X/46,XX. American journal of medical genetics Part A 2016; 170a(5): 1196–201.

3. Viuff MH, Stochholm K, Uldbjerg N, Nielsen BB, Gravholt CH. Only a minority of sex chromosome abnormalities are detected by a national prenatal screening program for Down syndrome. Human reproduction (Oxford, England) 2015; 30(10): 2419–26.

4. Koeberl DD, McGillivray B, Sybert VP. Prenatal diagnosis of 45,X/46,XX mosaicism and 45,X: implications for postnatal outcome. American journal of human genetics 1995; 57(3): 661–6.

5. Murdock DR, Donovan FX, Chandrasekharappa SC, et al. Whole-Exome Sequencing for Diagnosis of Turner Syndrome: Toward Next-Generation Sequencing and Newborn Screening. The Journal of clinical endocrinology and metabolism 2017; 102(5): 1529–37.

6. Wigby K, D’Epagnier C, Howell S, et al. Expanding the phenotype of Triple X syndrome: A comparison of prenatal versus postnatal diagnosis. American journal of medical genetics Part A 2016; 170(11): 2870–81.

7. Bucerzan S, Miclea D, Popp R, et al. Clinical and genetic characteristics in a group of 45 patients with Turner syndrome (monocentric study). Therapeutics and clinical risk management 2017; 13: 613–22.

8. Lopez L, Arheart KL, Colan SD, et al. Turner syndrome is an independent risk factor for aortic dilation in the young. Pediatrics 2008; 121(6): e1622–7.

9. Morimoto N, Tanaka T, Taiji H, et al. Hearing loss in Turner syndrome. The Journal of pediatrics 2006; 149(5): 697–701.

10. Klaskova E, Zapletalova J, Kapralova S, et al. Increased prevalence of bicuspid aortic valve in Turner syndrome links with karyotype: the crucial importance of detailed cardiovascular screening. Journal of pediatric endocrinology & metabolism: JPEM 2017; 30(3): 319–25.

11. Abd-Elmoniem KZ, Bakalov VK, Matta JR, et al. X chromosome parental origin and aortic stiffness in turner syndrome. Clinical endocrinology 2014; 81(3): 467–70.

12. Folsom LJ, Fuqua JS. Reproductive Issues in Women with Turner Syndrome. Endocrinology and metabolism clinics of North America 2015; 44(4): 723–37.

13. Stochholm K, Juul S, Juel K, Naeraa RW, Gravholt CH. Prevalence, incidence, diagnostic delay, and mortality in Turner syndrome. The Journal of clinical endocrinology and metabolism 2006; 91(10): 3897–902.

14. Russell LM, Strike P, Browne CE, Jacobs PA. X chromosome loss and ageing. Cytogenetic and genome research 2007; 116(3): 181–5.

15. Sudlow C, Gallacher J, Allen N, et al. UK biobank: an open access resource for identifying the causes of a wide range of complex diseases of middle and old age. PLoS medicine 2015; 12(3): e1001779.

16. Tukey JW. Exploratory data analysis. Reading, Mass.: Addison-Wesley Pub. Co.; 1977.

17. Tartaglia NR, Howell S, Sutherland A, Wilson R, Wilson L. A review of trisomy X (47,XXX). Orphanet journal of rare diseases 2010; 5: 8.

18. Cameron-Pimblett A, La Rosa C, King TFJ. The Turner syndrome life course project: Karyotype-phenotype analyses across the lifespan. 2017.

19. Fry A, Littlejohns TJ, Sudlow C, et al. Comparison of Sociodemographic and Health-Related Characteristics of UK Biobank Participants with the General Population. American journal of epidemiology 2017.

20. Gravholt CH, Andersen NH, Conway GS, et al. Clinical practice guidelines for the care of girls and women with Turner syndrome: proceedings from the 2016 Cincinnati International Turner Syndrome Meeting. European journal of endocrinology 2017; 177(3): G1–g70.

21. Jacobs PA, Baikie AG, Brown WM, Macgregor TN, Maclean N, Harnden DG. Evidence for the existence of the human “super female”. Lancet 1959; 2(7100): 423–5.

22. Ottesen AM, Aksglaede L, Garn I, et al. Increased number of sex chromosomes affects height in a nonlinear fashion: a study of 305 patients with sex chromosome aneuploidy. American journal of medical genetics Part A 2010; 152a(5): 1206–12.

23. Leggett V, Jacobs P, Nation K, Scerif G, Bishop DV. Neurocognitive outcomes of individuals with a sex chromosome trisomy: XXX, XYY, or XXY: a systematic review. Developmental medicine and child neurology 2010; 52(2): 119–29.

24. Stochholm K, Juul S, Gravholt CH. Poor socio-economic status in 47,XXX —an unexpected effect of an extra X chromosome. European journal of medical genetics 2013; 56(6): 286–91.

25. Alexander M, Ding Y, Foskett N, Petri H, Wandel C, Khwaja O. Population prevalence of Down’s syndrome in the United Kingdom. Journal of intellectual disability research: JIDR 2016; 60(9): 874–8.

26. James RS, Dalton P, Gustashaw K, et al. Molecular characterization of isochromosomes of Xq. Annals of human genetics 1997; 61(Pt 6): 485–90.

